# Proteus: A Random Forest Classifier to Predict Disorder-to-Order Transitioning Binding Regions in Intrinsically Disordered Proteins

**DOI:** 10.1101/080788

**Authors:** Sankar Basu, Fredrik Söderquist, Björn Wallner

**Affiliations:** Bioinformatics Division, Department of Physics, Chemistry and Biology, Linköping University, Linköping, Sweden; Department of Biochemistry, University of Calcutta, Kolkata – 700019, India; Swedish e-Science Research Center, Linköping University, Sweden.

**Keywords:** Intrinsic Disorder, Protean, Random, Forest, Disorder-to-order Transition, Topography length

## Abstract

The focus of the computational structural biology community has taken a dramatic shift over the past one-and-a-half decades from the classical protein structure prediction problem to the possible understanding of intrinsically disordered proteins (IDP) or proteins containing regions of disorder (IDPR). The current interest lies in the unraveling of a disorder-to-order transitioning code embedded in the amino acid sequences of IDPs / IDPRs. Disordered proteins are characterized by an enormous amount of structural plasticity which makes them promiscuous in binding to different partners, multi-functional in cellular activity and atypical in folding energy landscapes resembling partially folded molten globules. Also, their involvement in several deadly human diseases (e.g. cancer, cardiovascular and neurodegenerative diseases) makes them attractive drug targets, and important for a biochemical understanding of the disease(s). The study of the structural ensemble of IDPs is rather difficult, in particular for transient interactions. When bound to a structured partner, an IDPR adapts an ordered conformation in the complex. The residues that undergo this disorder-to-order transition are called protean residues, generally found in short contiguous stretches and the first step in understanding the modus operandi of an IDP / IDPR would be to predict these residues. There are a few available methods which predict these protean segments from their amino acid sequences; however, their performance reported in the literature leaves clear room for improvement. With this background, the current study presents 'Proteus', a random forest classifier that predicts the likelihood of a residue undergoing a disorder-to-order transition upon binding to a potential partner protein. The prediction is based on features that can be calculated using the amino acid sequence alone. Proteus compares favorably with existing methods predicting twice as many true positives as the second best method (55% vs. 27%) with a much higher precision on an independent data set. The current study also sheds some light on a possible 'disorder-to-order' transitioning consensus, untangled, yet embedded in the amino acid sequence of IDPs. Some guidelines have also been suggested for proceeding with a real-life structural modeling involving an IDPR using Proteus.

**Software Availability:** https://github.com/bjornwallner/proteus

## Introduction

After extensive research over one-and-a-half decades, it is evident that many functional proteins lack well-folded 3D structures. These intrinsically disordered proteins (IDPs), could be completely disordered or contain intrinsically disordered protein regions (IDPRs) [1–5]. In contrast to the classical view of protein folding [6], where a nascent cytoplasmic polypeptide chain folds into a stable globule, concomitantly while being synthesized [7,8], these proteins are born disordered [3] and remain either completely or partially unstructured throughout their entire life span. It is only when they interact with functionally relevant binding partners that they switch to ordered structures [4]. In fact, their existence in a biologically active form without adapting to a unique 3D-structure contradicts the traditional notion of the “one protein–one structure–one function” paradigm [1].

IDPs are highly abundant in nature and have been found to be involved in a number of functions within the living cell, most of which belong to the non-classic (non-enzyme) type [9,10]. They possess remarkable binding promiscuity [4] in a wide range of intermolecular interactions, complementing the functional repertoire of ordered globular proteins, similar to the phenomena of enthalpy – entropy compensation [11]. The promiscuity is primarily manifested in their ability to interact specifically with structurally diverse molecular partners and obtaining different structures upon binding. It is highly likely that these peculiar characteristics may be attributed to their nonnative-like <u>multi-funneled</u> and relatively flat energy landscapes [12,13], wherein the favored conformations closely resemble to the partially folded molten globules [13] which also enable them to preserve the necessary amount of disorder even in their bound forms [4]. Considering this flexible nature, they have been referred to as part of the 'edge of chaos' systems [14], serving as a bridge between well-ordered and chaotic system that is critical in the context of cellular energy balance.

In addition to these peculiar biophysical and folding attributes, IDPs are also of considerable biomedical interest due to their functional importance. In fact, the functions they are involved in (e.g., regulation, signaling, and control) are mostly the ones that require high specificity – low-affinity interactions [15]. Recent studies have highlighted their multifarious activities as molecular rheostats and molecular clocks, in tissue specific and alternative splicing of mRNA, transport of rRNA and proteins and RNA-chaperons [16]. Also, by sustaining enough disorder even in the bound form, IDPs are enabled to participate in both one-to-many and many-to-one signaling [2]. The promiscuity in binding also suggests that not only misfolding [17], but also misidentification or mis-signaling [2] in biomolecular recognition could serve as the root cause of some extremely complex human diseases [3] including cancer, diabetes, amyloidoses, and cardiovascular and neurodegenerative diseases [18].

Taken together, there is a great need for a deeper understanding of IDPs and their interactors. However, since obtaining information about IDPs from experiments is difficult owing to their inherent disorder, computational modeling provides a realistic way forward. For most IDPs, only a subset of the disordered residues can actually undergo a disorder-to-order transitions upon binding to a folded protein, leading to the concept of ‘folding coupled with binding’ [19]. These segments are called *protean* borrowed from Greek mythology, meaning ‘ever-changeable’ or ‘mutable’ [19]. To model the 3D structure of an interacting IDP / IDPR, a first aim would be to predict the potential ‘mutable’ *protean* regions. It is important to note that, due to the intrinsic disorder, these regions in an isolated X-ray structure are presented as ‘missing electron density’ patches (listed in REMARK 465 in the corresponding PDB file [5]), and should only appear structured in its bound form. This is in fact also the definition of a ‘protean’ segments.

Extensive studies have analyzed the sequence space of IDPs in relation to their intrinsic disorder. These studies reveal their correspondence to low entropy sequences with less complexity [5,20]. In particular, tandem repeats have often been found to be embedded in these sequences (e.g. polyglutamine stretches in amyloid beta [21]) giving rise to the notion of ‘the more perfect the less structured’ proteins [22]. Thus, in a sense, low sequence entropy can potentially lead to high conformational entropy, characteristic of the IDPs. Some mechanistic insights into the origin of the disorder have also been suggested, for example, the low content of hydrophobic residues with an abundance of charged residues in IDPs [23] disfavoring self-folding [24] by potentially decreasing the number of possible two-body contacts [25]. Furthermore, the charge – hydrophobicity boundary have been envisaged to represent a trade-off between repulsive and attractive interactions reminiscent of globular – disorder transitions [26].

Nevertheless, it remains highly challenging to decipher the root cause of intrinsic disorder from pure sequence-based investigation given the limited structural data. Concerted efforts have been made to untangle a possible disorder code from amino acid sequence alone which includes deciphering the propensity for intrinsic disorder [26], and statistical mechanical potentials describing sequence-derived elasticity [27]. The nature of the problem is ideal for machine learning algorithms given the availability of annotated sequence data. In fact, quite a few predictors have recently been developed that predict not only the disordered regions [28–32], but also the ‘protean’ segments [32–35]. Still, ‘protean prediction’ is in an early stage, offering much room for improvement. In this background, the current study does not only attempt to shed some light on a possible yet unexplored sequence consensus of such ‘disorder-to-order’ transitions, but also presents ‘Proteus’, a random forest classifier that predicts protean segments solely from the amino acid sequence of an IDP. Proteus compares favorably to the existing predictors. Some guidelines have also been suggested on how and where to use Proteus during the course of a real-life structural modeling involving an IDPR.

## Methods

### Training Dataset

Two databases containing proteins with annotated protean segments, IDEAL[19] and MoRF[34] were pulled together to build the final training dataset. IDEAL (Intrinsically Disordered proteins with Extensive Annotations and Literature) contains 557 proteins with experimentally verified protean segments called ‘ProS’ in the database. However, only 203 of 557 proteins in this database actually contain protean segments. The rest are IDPs where no protean segments have yet been experimentally verified and thus serve as negative examples in training. The MoRF dataset comes from MoRFpred [34], one of the existing classifiers. It contains 840 proteins, and all of them have at least one protean segment. More importantly, all members of MoRF have direct structural evidence from the PDB. Members from IDEAL and MoRF will henceforth be referred to as ‘ProS’ and ‘MoRF’ respectively, and the combined dataset as ‘PnM’. The details of all datasets have been enlisted in Table 1.

**Table 1.** Description of the datasets.

| Dataset | Proteins | Protean Residues | Non-Protean Residues | Total Residues |
| --- | --- | --- | --- | --- |
| ProS | 557 | 6,245 | 356,053 | 362,298 |
| MoRF | 840 | 10,549 | 494,264 | 504,813 |
| ProS + MoRF (PnM) | 1,397 | 16,794 | 850,317 | 867,111 |
| Validation | 9 | 163 | 2,046 | 2,209 |

### Independent Benchmark

Nine proteins that were used as independent benchmark in the DISOPRED3 study [32] were used as an independent benchmark set here as well. In the DISOPRED3 study 29 chains having protean segments were initially culled using database annotations and publications, later they had to be reduced to nine proteins, as the other 20 chains were found to be used in the training datasets of the competing methods, ANCHOR [33], MoRFpred [34] and MFSPSSMPred [35]. None of the nine proteins were similar to any protein in the current training dataset.

### Target Function

The binary status for each amino acid residue in the sequence to be protean or non-protean was used as the target function in training the classifier, denoted by 1 and 0 respectively, for positive and negative examples. It is to be noted that non-protean residues refer to the ordered as well as disordered residues which do not undergo the ‘disorder-to-order transition’ upon binding and hence, remain disordered even in the bound state.

### Data Clustering and Cross-Validation Benchmark

To avoid training and testing on similar examples, BLASTclust was used to cluster the protein sequences in the combined dataset ‘PnM’. Sequences with a pairwise similarity of at least 30% over at least 50% of the sequence length (-S 30 -L 0.5) were clustered. This resulted in 774 clusters, the largest containing 38 proteins, and 253 clusters containing more than one protein. One third of all ProS sequences were found to be similar to at least one MoRF sequence and vice-versa.

To prepare the data for five-fold cross-validation, five folds were built by grouping clusters in such a way that the number of target proteins remain consistent among the folds. This resulted in four folds with 280 targets and one fold with 279 targets, containing between 158,651 and 218,870 amino acid residues, and around 1.4% to 2.2% positive examples. During cross-validated training, four of the folds are used for training and the remaining one is used for testing. This is repeated five times to make predictions for all five folds.

### Random Forest Classifier

The random forest classifier module in scikit-learn Python package [36] was used for training. Every decision tree in the forest classify examples as positive or negative, and a final decision is made according to a majority vote.

### Evaluation Measures

In binary classification, there are four possible outcomes when classifying an example: (i) True Positive (TP): a positive example correctly classified as positive; (ii) True Negative (TN): a negative example, correctly classified as negative; (iii) False Negative (FN): a positive example incorrectly classified as negative; and (iv) False Positive (FP): a negative example incorrectly classified as positive. By counting these four possible outcomes, the following evaluation measures were calculated.

#### Precision (PPV)

Precision, also known as specificity or the Positive Predicted Value (PPV), measures how many examples classified as positive were actually positive, calculated by the ratio, TP / (TP + FP).

#### Recall (TPR)

Recall (or coverage) measures how many positive examples were correctly classified as positives. It is also called the ‘True Positive Rate’ (TPR) and calculated by the ratio, TP / ∑P, where ∑P is the total number of positives, i.e., ∑P = TP + FN.

#### F1-score

F1-score is the harmonic mean between PPV and TPR and could be interpreted as a trade-off between PPV and TPR. It is defined by the following equation: F1=2PPV×TPR/(PPV+TPR).

#### Matthews Correlation Coefficient

Another direct evaluation measure of classification performance is the Matthews Correlation Coefficient (MCC) ranging from −1 (perfect inverse prediction) to +1 (perfect prediction) and calculated as: MCC=((TP×TN)−(FP×FN))/((TP+FP)(TP+FN)(TN+FP)(TN+FN))^1/2^. This was used in conjugation with the F1-score to estimate the overall performance of the predictor.

### Tuning Training Parameters

#### Decision tree depth

In general the deeper the tree, the more complex patterns it can fit. However, this can easily lead to over-fitting. Thus, finding an optimal tree depth is important. The maximum depth was varied between 1 and 25 (**Supplementary Fig. S1**) and a depth of 13 yielded the highest MCC and F1 scores.

#### Number of trees in the forest

Another important parameter is how many decision trees to use. In theory, the more trees the better, but there is a saturation in performance, beyond which the increase in performance is only marginal. Therefore, it is important to find the optimal number of trees to save computational time. As can be seen from the **Supplementary Fig. S2**, 50 decision trees yield a reasonable performance, which is only slightly increased (by ∼5%) using more trees. Therefore, using 50 trees was considered to be enough for the computationally expensive feature selection part. However, for the final selected combination of features, 500 trees were used to achieve maximum performance.

#### Probability cutoff

The classifier needs a user-defined probability cutoff (P_cut_) above which an example is classified as positive. P_cut_ was varied in the whole range of 0.0 to 1.0 and based on the performance (**Supplementary Fig. S3**), was set to 0.5 (majority vote). Therefore, if 50% or more decision trees voted for the particular example to be positive, it was classified as positive.

### Frequency Analyses of Protean and Disordered Residues

#### Amino Acid Propensity

The propensity (Pr) for a particular amino acid, X to occupy a particular ‘class’ (e.g. protean vs. disordered residues) was calculated as the ratio of two probabilities (P) as: Pr(X) = P(X)_class_/P(X)_full_ = (N(X)_class_/N(All)_full_) / (N(X)_full_ / N(All)_full_) where ‘full’ stands for the entire training dataset and N denotes the raw count of amino acid(s) in the said ‘class’. A propensity value of 1 represents no preference whereas a higher or lower value represent higher or lower preference, respectively, of the amino acid to occupy the given class with respect to the baseline.

#### Predicted Secondary Structural Content

PSIPRED [37] was used to predict the secondary structure in three classes (H: Helix, E: Strand, C: Coil). For each amino acid, the relative fraction of each of the three main secondary structural classes (H, E, C) were calculated for protean, non-protean, disordered and ordered sequences. The aim was to decipher if there was any preference in disorder vs. order sequences that might have propagated to protean segments during the ‘disorder-to-order’ transitions.

### Design of the sequence-driven features

#### Consideration of local and global effects

The origin of disorder is a conjunction of multiple factors. Therefore, ideally the contribution of both, the local sequence (neighboring effect) and that of the global three-dimensional fold of the protein should be considered in the design of features. However, it is highly non-trivial to take into account the global effect of the overall protein fold without actually attempting to build homology models for the predicted ‘structured’ regions, in their bound form. Incorporating such a modeling pipeline will be computationally costly and will also have low confidence associated with the built models due to the lack of enough structural data. One alternative way to indirectly take into account the global constraints is to perform a homology search against all non-redundant sequences [38] and then convert the sequence into a profile. To this end, PSI-BLAST [39] was used to construct sequence profiles. In addition, PSIPRED [37] was used to predict secondary structure and DISOPRED3 [32] was used to predict disorder probability for each amino acid residue. Thus, the plausible global constraints were also accounted for in the designed features, at least implicitly.

To describe the neighboring environment, a sliding window of 15-residues centered around the current residue was considered in the design of most features. This will produce an average property of the feature, taking into account the local sequence dependence associated with disorder-to-order transitions. The size of the window was optimized by trying different lengths in the range of 9-21. The optimal size agrees with the average length of protean segments (Fig.1).

**Fig.1.**
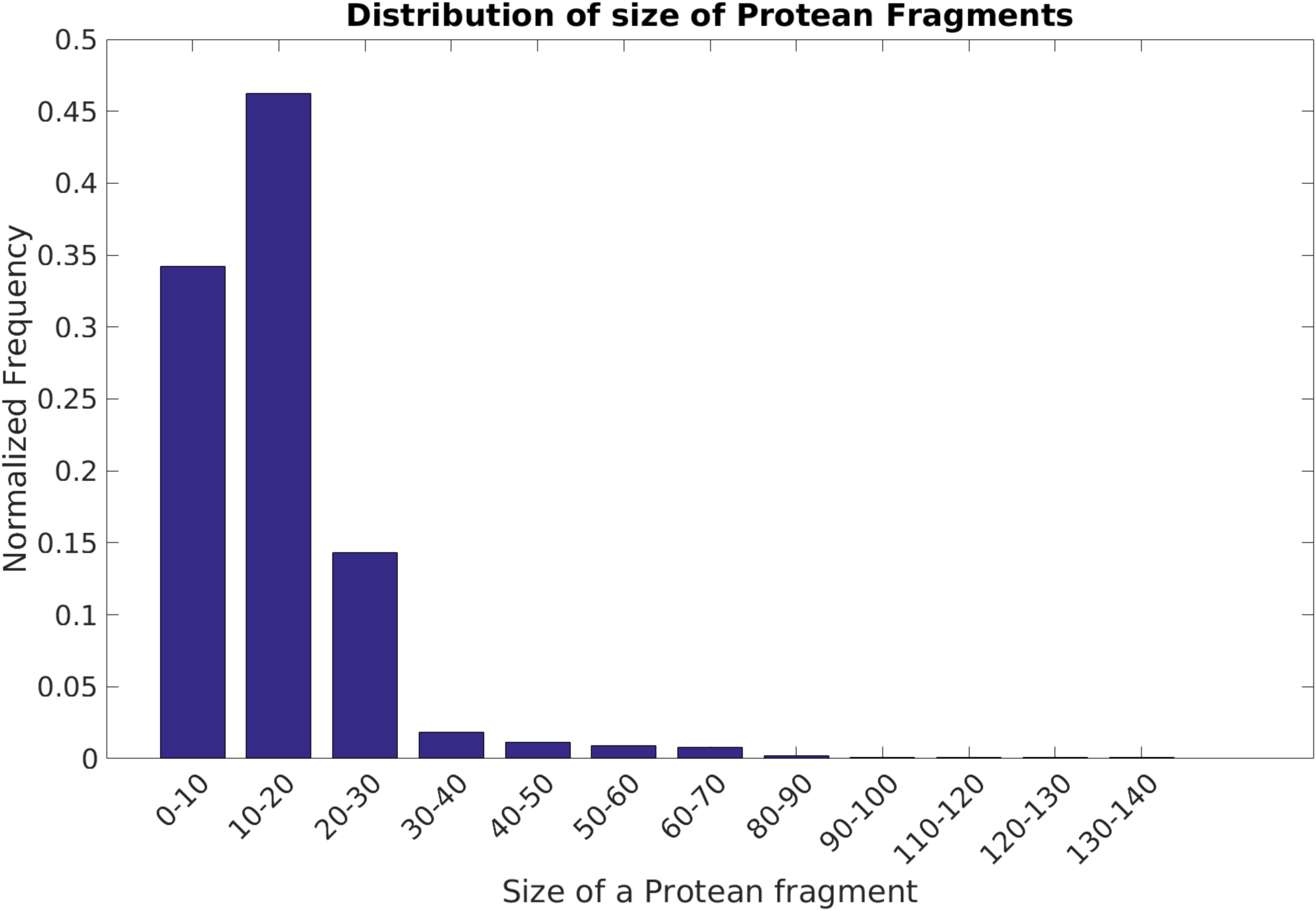
Distribution of size of the ‘annotated’ protean segments. The distribution is obtained from the combined ‘PnM’ training dataset.

In total 342 features, in seven different feature groups, were used and are described in detail below (Table 2)

**Table 2.** A Summary of Feature groups.

| Feature Group | Name | Feature Number | Count |
| --- | --- | --- | --- |
| 1 | Sequence Profile | 1-300 | $20 \times 15 = 300$ |
| 2 | Amino Acid Conservation | 301 | 1 |
| 3 | Amino Acid Concentration | 302-321 | $20 \times 1 = 20$ |
| 4 | Amino Acid Properties | 322-330 | $4 + 3 + 1 + 1 = 9$ |
| 5 | Predicted Secondary Structure | 331-333 | $3 \times 1 = 3$ |
| 6 | Predicted Disorder | 334-340 | $3 + 3 + 1 = 7$ |
| 7 | Disorder Topography | 341-342 | $1 + 1 = 2$ |

#### Feature Group 1: Amino Acid Mutability (Features: 1-300)

Considering the influence of the local sequence on disorder, it is likely that empirical trends (over and under-representations) will be found in the distribution of amino acids in protean compared to non-protean regions. In other words, certain amino acids might preferentially occur in the protean segments but not others. This was represented by Position Specific Scoring Matrices (PSSM) constructed by running three iteration (-j 3) of PSI-BLAST [39] against UniRef90 [38] with an inclusion E-value threshold of 10^−3^ (-h 0.001). The PSSM contains scores for each of the 20 possible amino acid substitutions in each position, representing the amino acid mutability at any given position. The higher the score, the higher the probability that these amino acids occurs at that position. To improve convergence, the raw PSSM scores were linearly scaled to [0.0, 1.0] based on the maximum and minimum values observed for each amino acid in the whole training set. To account for the local sequence bias, a 15-residue window of the PSSM was used centered around the current residue, giving 300 (15×20) features in total for each residue.

#### Feature Group 2: Amino Acid Conservation (Feature: 301)

The conservation score is derived by PSI-BLAST [39] from the PSSM matrix, and, as the name suggests, conceptually, it is complementary to that of ‘mutability’. Numerically, it is a modified Shannon Entropy [40] term representative of the heterogeneity of amino acid substitutions for a given position in the input sequence. Again, to take care of the neighboring environment, the conservation score was averaged over a 15-residue window. In contrast to all other feature groups, this group consists of only a single value.

#### Feature Group 3: Amino Acid Composition (Features: 302-321)

This feature group describes the individual concentration of all amino acids, in a 15-residue long window, i.e. 20 features in all, representing a coarse-grain estimation of the amino acid properties in the local neighborhood around the central residue.

#### Feature Group 4: Amino Acid Properties (Features: 322-330)

It is natural to believe that the physiochemical properties of different amino acids hold the key for developing intrinsic disorder and also for the disorder-to-order transitions. In contrast to the ‘amino acid composition group’ described above, Polarity, Charge, Hydrophobicity and Molecular Weight were explicitly described in this feature-group, in a 15-residue sliding window. Polarity was divided into polar, non-polar, acidic-polar or basic-polar, and charge into positive, negative, and neutral [41]. Hydrophobicity was described using the Kyte Doolittle scale [42]. For each of these seven features, the corresponding counts were averaged over the 15-residue window.

#### Feature Group 5: Predicted Secondary Structure (Features: 331-333)

Secondary structural propensities of individual amino acids in the close neighborhood of a residue might have major influence on disorder and might serve as a discriminative feature between protean and non-protean fragments. For example, if this likelihood keeps altering between helices to sheets along the sequence, the resultant main-chain trajectory would potentially keep wobbling giving rise to an unstructured region. The other possibility is of course having most residues predicted as ‘random coils’. The probabilities of each amino acid residue in a sequence to form one of the three main secondary structures (Helix, Strand, Coil) were predicted by PSIPRED[37] and averaged over a 15-residue sliding window, serving as three distinct features.

#### Feature Group 6: Predicted Disorder Probability (Features: 334-340)

The probability for disorder was predicted using DISOPRED3 [32]. The disorder prediction score from DISOPRED3 is a confidence estimate (or probability) for a residue in a protein sequence to be disordered. It is defined in the range [0, 1] and DISOPRED3 assigns the disordered status to a residue if the score is greater than 0.5. The disorder prediction score, averaged over the 15-residue window centered on the current residue was directly used as the first feature in this group. In addition, to describe the local properties of the disorder prediction, the length of disordered and ordered segments and the start and end positions relative to the total sequence length were also used. In detail, if the score was greater than 0.5, the positions on either side of the current residue where the score drops below 0.5 were identified. From this, the length, start and stop positions of the segment could be calculated. This was performed for residues predicted to be disordered (score > 0.5) and for residues predicted to be ordered (score < 0.5), resulting in 7 (1+3+3) features. Depending on the predicted disorder of the segment, three of the seven features will always remain zero.

#### Feature Group 7: Disorder Topography (Features: 341-342)

Disorder topography measures the topography of peaks and valleys in the predicted disorder score graph (**Supplementary Fig. S4**). Each residue is classified as being part of a peak (1), valley (-1) or neither (0). A residue is part of a peak if on both sides, there exists another residue with a score at least 10% lower than the current residue. Likewise, a residue is part of a valley if there are residues with disorder scores at least 10% higher than the current residue. If a residue is neither at a peak nor in a valley it is classified as neither. In addition, the length of the current peak or valley residue is also calculated and used as a separate feature. Thus, the disorder topography feature consists of the peak/valley/neither classification (feature no. 341) and the topographic length of the current peak/valley (feature no. 342).

## Results and Discussion

### Propagation of sequence consensus during disorder-to-order transitions

Sequence-driven properties such as amino acid propensities and predicted secondary structural content might serve as crucial consensus in the information transfer during the ‘disorder-to-order’ transition. A comparative study of these properties in predicted disordered and annotated protean segments will also serve to explore and identify empirical trends in the designed features and thereby act as a guide in determining the features that are more discriminative compared to the features that can act as filters. Taking this into account, the referred properties were investigated in (i) protean vs. non-protean residues and (ii) disordered vs. ordered residues (as predicted by DISOPRED3 [32]) and compared with each other. The aim was to identify any pattern that might be responsible for the disorder-to-order transitions, implicitly embedded in the protean sequences. To that end, we wanted to collect the most discriminating trends in the disordered vs. ordered regions which were either maintained or inverted in the protean vs. non-protean segments. These combined trends should be instrumental in both sustaining the intrinsic disorder and also in the information transfer during the ‘disorder-to-order’ transitions. However, since the ‘disorder vs. order’ classification is clearer and more distinct, it was expected that the trends for ‘disorder vs. order’ should be more prominent than the ‘protean vs. non-protean’ trends.

### Amino acids preference in protean and disorder residues

The first and most fundamental characteristic investigated was the of amino acid propensity in disordered, ordered and protean / non-protean residues. The predicted disordered regions show drastic under-representations of hydrophobic amino acids compared to predicted ordered regions (Fig. 2A). Even among the distribution of hydrophobic amino acids, there is an unmistakable trend with respect to the size of the hydrophobic side-chain. The gradual increase in the propensity of the hydrophobic side-chains in the predicted ordered regions is found to be directly proportional to their side-chain volume (Ala → Val → Leu → Ile → Phe → Tyr → Trp) (Fig. 2B); whereas in the predicted disordered regions, the relationship appears inversely proportional. This trend is perfectly consistent with the notion of hydrophobic core formation within ordered protein tertiary structures [43], and on the other hand, bulky aromatics (Phe, Tyr, Trp) should be unfavorable in disordered regions, due to their potential incompatibility with regard to side-chain volume and entropy. The other noticeable features are the significant over-representation of cysteines in ordered regions with a concomitant under-representation in disordered regions, consistent with the idea of fold stabilization by disulfide bridges [44], which must be avoided during the natural design of intrinsic disorder. On the other hand, prolines are significantly over-represented in disordered compared to ordered regions, which is consistent with their ability to break regular secondary structures [45], especially helices [46]. Even if found in regular secondary structures (β-sheets for example), proline needs additional structural constraints from pre-prolines (e.g., glycine rescue) to become stabilized [47]. In line with these observations, proline has been identified as the most disorder promoting amino acid residue [48].

**Fig.2.**
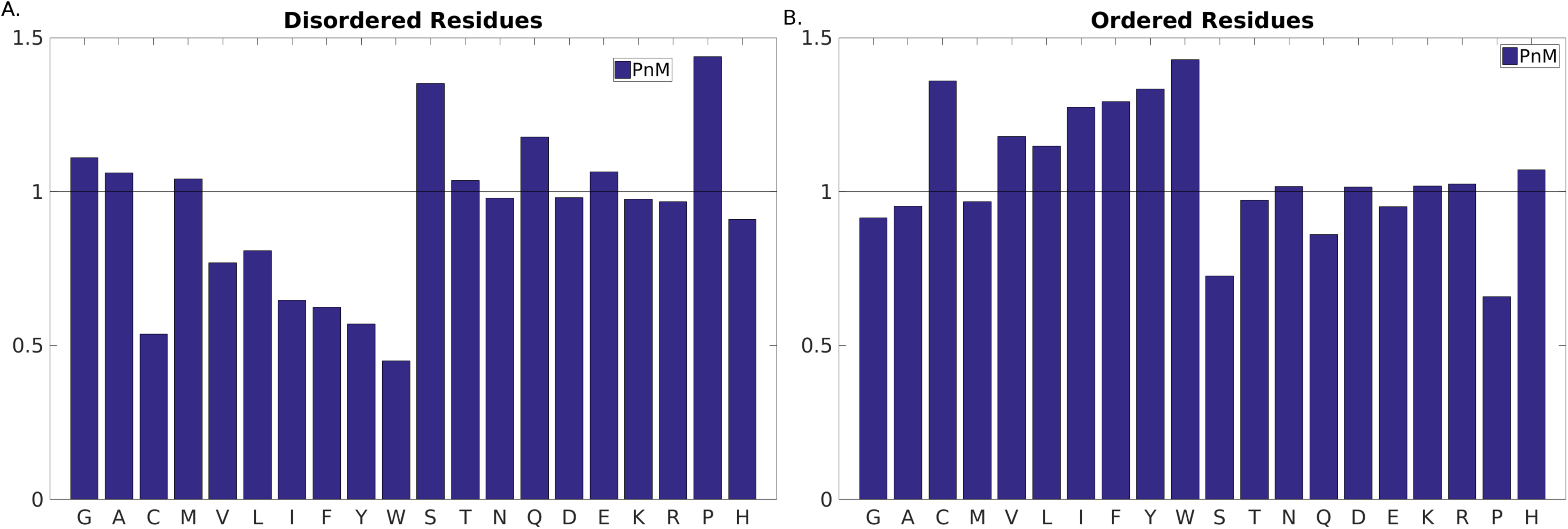
Amino Acid Propensities in the ‘predicted’ disordered vs. ordered regions. The Black Horizontal Line (Propensity = 1.0) serves as the baseline; meaning no preferential occurrence of the said amino acid in the said class. A propensity greater or lesser than 1.0 represents over and under representations respectively.

The other well-known residue, responsible for backbone flexibility, glycine [45] was also found to be over-represented in disordered compared to ordered regions. This is in accord with the well-established idea that prolines and glycines are general indicators of entropic elasticity [27,48] and hence control self-organization of elastomeric proteins (e.g., amyloid fibrils) [49]. In fact, recent studies have formulated correlation functions of elasticity in terms of coiling propensity based on sequences rich in proline and glycine in disordered proteins [27,48].

The other noticeable difference was seen for serine, again a small and polar amino acid, significantly over-represented in disordered and under-represented in ordered regions. Indeed, serine-rich proteins in bacterial enzymes like kinases [50] and eukaryotic splicing factors [51] have been reported to be part of intrinsically disordered proteins. The other polar (Thr, Asn, Gln) and charged (Asp, Glu, Lys, Arg) amino acids were found to have similar or slightly higher propensities in disordered compared to ordered sequences, which agrees well with the earlier observations [48] [47].

But as mentioned earlier, the focus of the current work was to identify patterns that were not only discriminative in disorder vs. order sequences but were also maintained in protean vs. non-protean sequences and therefore might help in establishing a crucial consensus in the understanding of disorder-to-order transitions. However as expected, the patterns in protean vs. non-protean sequences were not as prominent as in disordered vs. ordered sequences (Fig. 3). The collection of all non-ProS plus non-MoRF sequences served as the ‘non-protean’ baseline which raised a value of ∼1.00 (+/- 0.01) for the baseline propensities of all amino acids (Fig. 3B). This was not surprising since the bulk majority of the training dataset contained negative examples (non-protean sequences). Similar to amino acid propensities obtained for the ordered regions, all large hydrophobic residues (Leu, Ile, Phe, Tyr, Trp) were found to be over-represented in the protean segments (Fig. 3A) and drastically under-represented in the disordered regions (Fig. 2A). This inversion in trends from disordered to protean segments is rather interesting since the protean segments are merely subsets of the originally disordered regions. It strongly indicates that the potential to get ordered by mediating enough hydrophobic interactions is in fact implicitly embedded in the protean sequences, just like that of globular proteins, but masked by neighboring or flanking disordered residues in their unbound forms.

**Fig.3.**
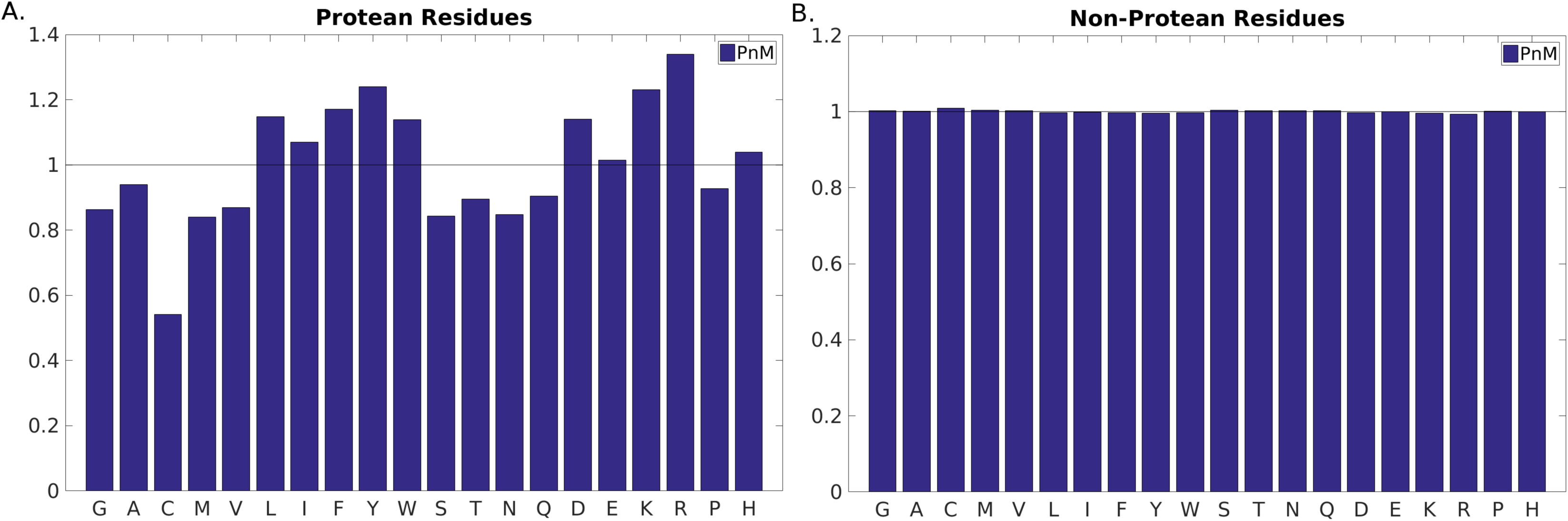
Amino Acid Propensities in the ‘annotated’ protean vs. non-protean segments. The Black Horizontal Line (Propensity = 1.0) serves as the baseline; meaning no preferential occurrence of the said amino acid in the said class. A propensity greater and lesser or 1.0 represents over and under representations respectively.

At the same time, the charged charged residues (Glu, Asp, Lys, Arg) also acquired much larger propensities compared to what they had in disordered sequences, and also noticeably higher propensities compared to ordered sequences in general (Fig. 2B and Fig. 3A). The results clearly indicate that both large, hydrophobic and charged residues are preferentially selected during the ‘disorder-to-order’ transitions (via binding). In other words, not all disordered regions undergo the same transition; rather, there is a preferential selection of sequences containing large hydrophobic and charged residues leading to stabilization through hydrophobic and salt-bridge interactions at the protein-protein interface. This is in accord with the general notion of stability upon binding in protein-protein interfaces where both shape and electrostatic complementaries are crucial for binding [52,53].

Finally, as for disorder residues cysteines are clearly under-represented in protean residues as well, reflecting the fact that the stability of protean residues should not involve disulfide bridges (at the cost of massive loss of plasticity). However, in contrast to disordered residues both proline and glycine are under-represented in protean residues, indicating that these residues do not undergo disorder-to-order transition; instead, they remain disordered.

### Secondary structure preference in protean and disordered residues

It is also important to conceptualize the secondary structural trends during the course of disorder-to-order transitions. The relative content of coil (C), including loops and turns is higher than helix (H) and strands (E) in all classes of sequences ranging from disorder-to-order and from protean to non-protean. But when comparing between two opposite class (e.g. disordered vs. ordered), it is the relative increment in (H+E)/C that is of interest. On that note, ordered sequences naturally have far greater regular secondary structures (H+E) amounting to ∼50% of the whole population than disordered sequences (H+E: ∼15%; C: ∼85%) (Fig. 4). As expected, the relatively low proportion (∼15%) of helices and strands in disorder residues definitely increases the disorder-to-order transitions in protean segments (H+E:∼40%), which is roughly the same as in non-protean sequences (Fig. 5). Recall that the large majority of the non-protean sequences are in fact the usual ordered sequences and the subset of disordered sequences that gets ordered only constitute a small (leftover) fraction. Among the regular secondary structures, helices appear to be more prevalent in protean (∼32%) than non-protean segments (∼27%) whereas beta-strands seem to be slightly more preferred in non-protean (∼10%) compared to protean segments (∼5%).

**Fig.4.**
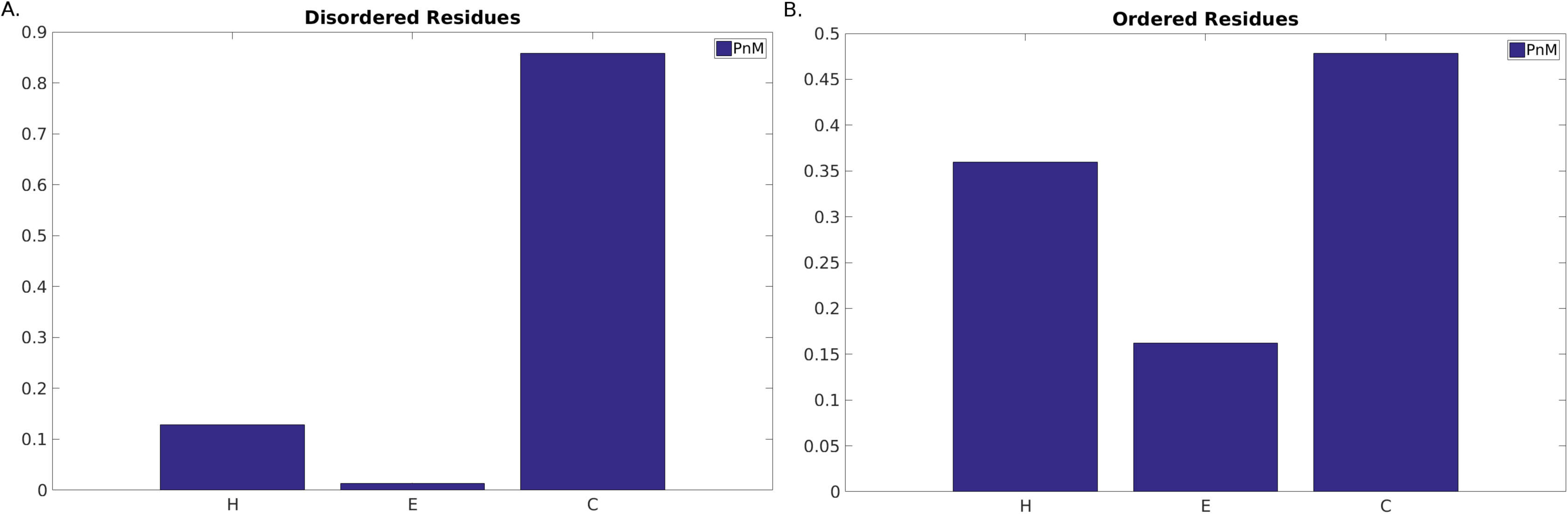
Secondary Structural probabilities in the ‘predicted’ disordered vs. ordered regions. H, E and C stands for α-Helix, β-Strand and Random Coil (non-helix, non-strand) respectively.

### Indecisiveness in adapting a particular secondary structure class from sequence

Another property investigated based on secondary structure is the indecisiveness of an amino acid sequence in adapting a particular secondary structure. The assumption being that protean segments, when disordered in isolation, might be indecisive in its choice to adapt a particular secondary structure (H, E or C) along their main-chain trajectory and thereby end up becoming unstructured. Given the current lack of structural data for these sequences, PSIPRED [37] was used to predict secondary structure and to test the above hypothesis. A measure for the indecisiveness or randomness in secondary structure prediction called Altscore was defined as the average number of transitions (H→C, C→E etc.) for each protean and non-protean segment. Regions with an Altscore value of ‘zero’ were omitted for both protean and non-protean regions, since they would only add noise to any potential signal. Focusing on the regions with Altscore> 0, the frequency distribution (Fig. 6) clearly discriminated between protean and non-protean classes with a wider spread being obtained for the protean class in addition to a peak-shift towards higher values (0.1 compared to 0.05 for non-protean). The results indicate that the intrinsic disorder associated with the unbound protean segments potentially suffers from the indecisiveness of the main-chain trajectory to adapt to a particular secondary structure.

**Fig.5.**
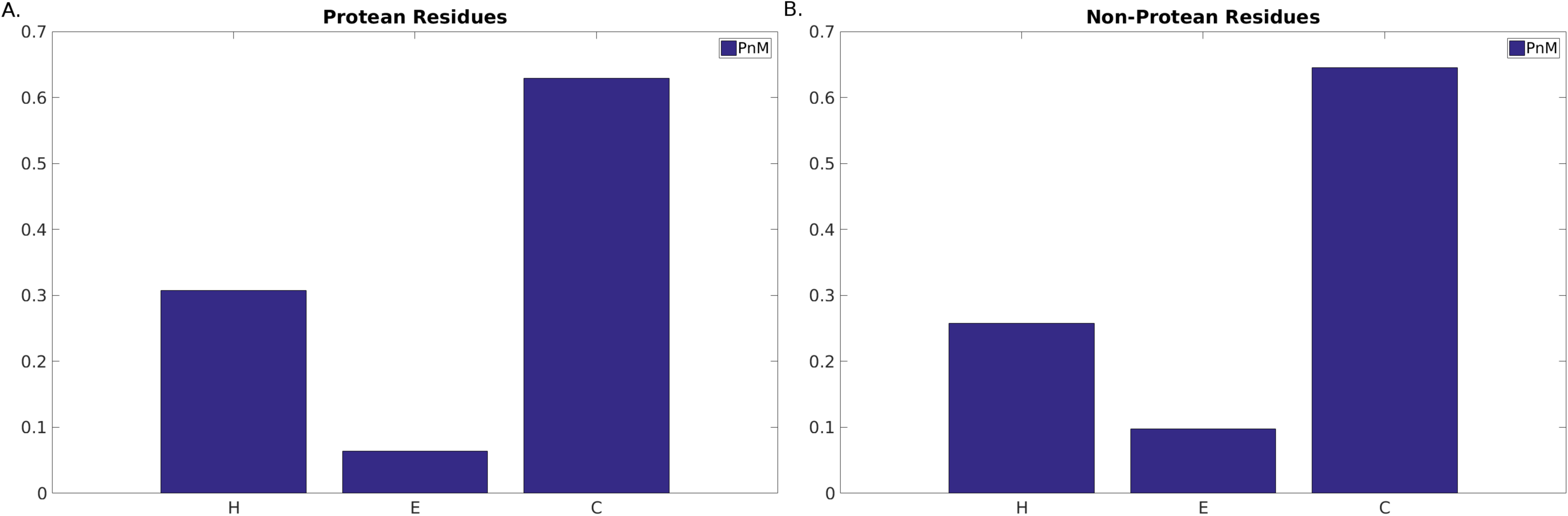
Secondary Structural probabilities in the ‘originally classified’ protean vs. non-protean segments. H, E and C stands for α-Helix, β-Strand and Random Coil (non-helix, non-strand) respectively.

**Fig.6.**
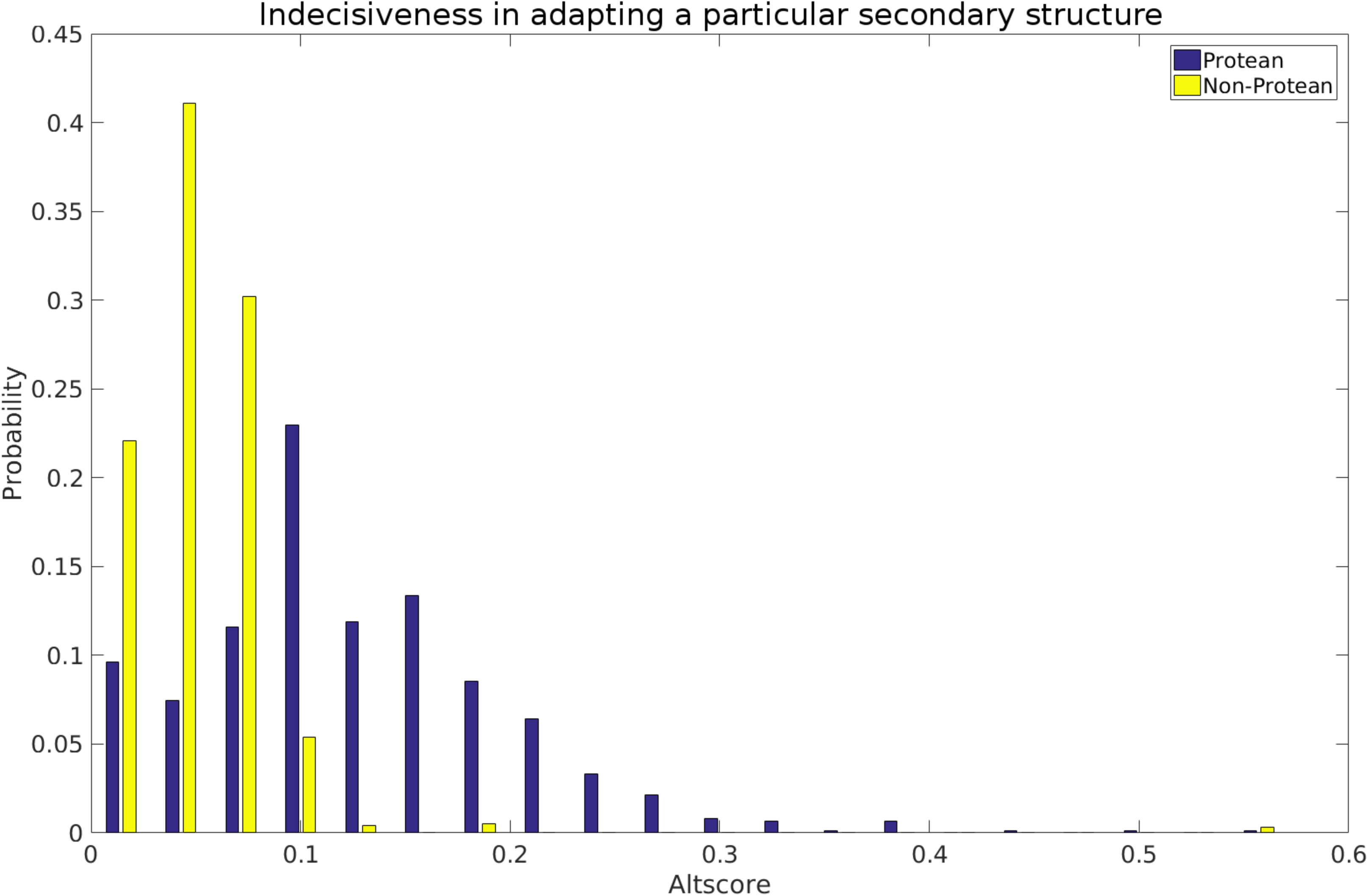
Indecisiveness in adapting a particular secondary structure for the ‘originally classified’ protean vs. non-protean segments. Probability Distributions of the Altscore (see Text) have been drawn for both sets. Segments assigned as purely ‘Coil’ were excluded from both sets.

Both the above observations, (i) the reappearance of large hydrophobic and charged amino acids into the protean segments, as well as (ii) the indecisiveness associated with their predicted secondary structures should serve constructively in unraveling a hidden consensus in promoting disorder-to-order transition.

### Training a classifier to predict protean residues

To be able to predict protean residues from sequence, a random forest classifier was trained on the features described in the Methods section. Most features were calculated using a 15-residue sliding, optimized by trying different window sizes in the range of 9-21 ( **Supplementary Fig. S5**). The chosen window size was optimal in the sense that it fell right in the center of the distribution of the protean segment-lengths (Fig. 1). An identical sliding window size was also used to determine protein-binding residues embedded within disordered regions previously [32]. Note that for all feature groups except Feature Group 1: Amino Acid Mutability, the number of features will remain the same even with a different window size. Among all features, some features might be non-informative, others might be redundant. Indeed, some features are similar in their physiochemical descriptions and therefore might be excluded without loss in performance. But sometimes it is advantageous for the classifier to learn from explicit rather than implicit features. To find the best combination of the seven feature groups, all 127 possible combinations were exhaustively examined by measuring the final cross-validated performance using MCC and F1-scores for each feature group combination.

The 20 best feature group combinations according to the MCC and F1-scores have been shown in **Supplementary Fig. S6** and **Fig. S7** respectively. The difference is small between the top feature group combinations. Also, the top-combinations as evaluated by MCC and F1 are not identical, whereas, using all features resulted in good scores being attained in both evaluations. Therefore, the combination of all feature groups was chosen judiciously. The absolute MCC and F1 score values are relatively small ∼0.13, owing to a large number of false positives and negatives. However, the magnitude of the scores are comparable to other studies [32–35], and reflect the difficulty of predicting residues that will be ordered upon binding from information in one of the binding partners only. This is further illustrated in the recall vs. precision (PPV) curves for the best combination (Fig. 7). The recall vs. precision curves were constructed by varying the cutoff (Pcut) and calculating precision and recall for each cutoff. The random base line precision is 1.9% and the curve for the best combination is clearly above that and it can also be seen that 500 trees is slightly better than 50. But the question remains of whether, the rather modest 10% precision at 23% recall (Pcut>0.5) is useful at all. Considering that it is still five times better than a random prediction, it is arguably useful given *the state-of-the-art*. But there is of course plenty of room for improvement which might be brought about in future studies by incorporating additional information not directly obtainable from the sequence alone. In principle, one might perform structure prediction with these sequences, and, during the course, filter out residues that are actually ordered by themselves; and also predict the surrounding residues. The predicted and demarcated structural units can also be used as starting templates in molecular dynamic simulations or docking studies exploring a reduced conformational space. In addition, advanced structural validation [54–56] could also be incorporated as filters in an iterative prediction pipeline to improve the sequence-based prediction.

**Fig.7.**
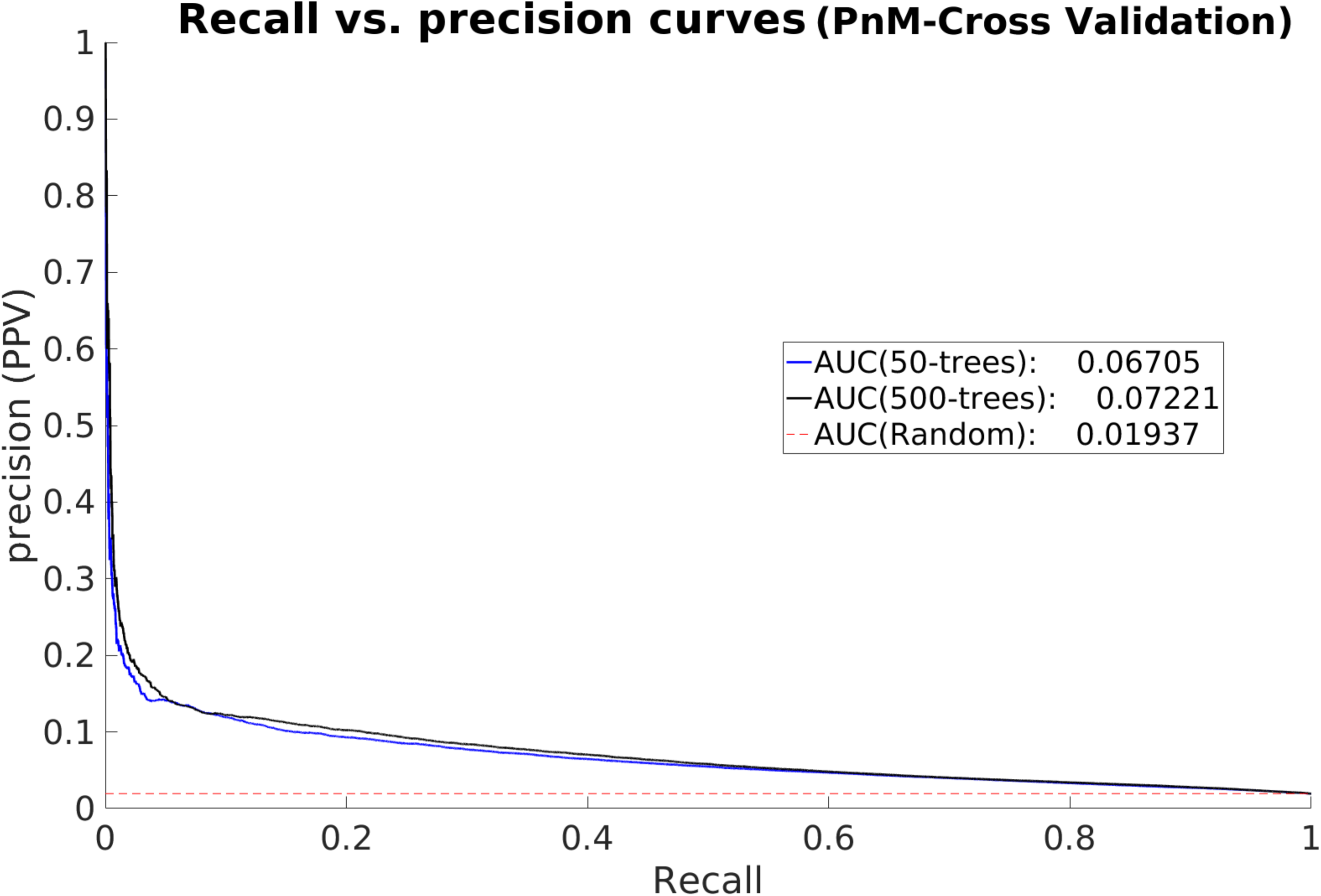
Recall vs. precision curves to analyze the cross-validated performance of Proteus. All five separate training / test folds as well as the final five-fold cross-validated ‘Proteus’ predictions (mean) are tabulated. The dashed line (- -) with a slope of 1.0 represents the random baseline.

### Relative Importance of Features

In an effort to learn what features contributed to the overall prediction, the relative importance of each feature group as calculated by the random forest prediction module was used. To take account of the inherent randomness associated with the classifications, this relative importance was averaged over predictions of 500 decision trees. As we can see, there are three features that stand out above the rest (Fig. 8): Topographic length (group 6) is by far the most important feature and describes the length of the topographic region where the current residue is located. Interestingly, the second most important feature is also a length descriptor, namely the more coarse-grained length of the ordered region corresponding to the current residue (group 5). Note that this feature will be ‘zero’ for all residues predicted to be disordered. The third most important feature is the predicted disorder score averaged over the current window size (group 5).

**Fig.8.**
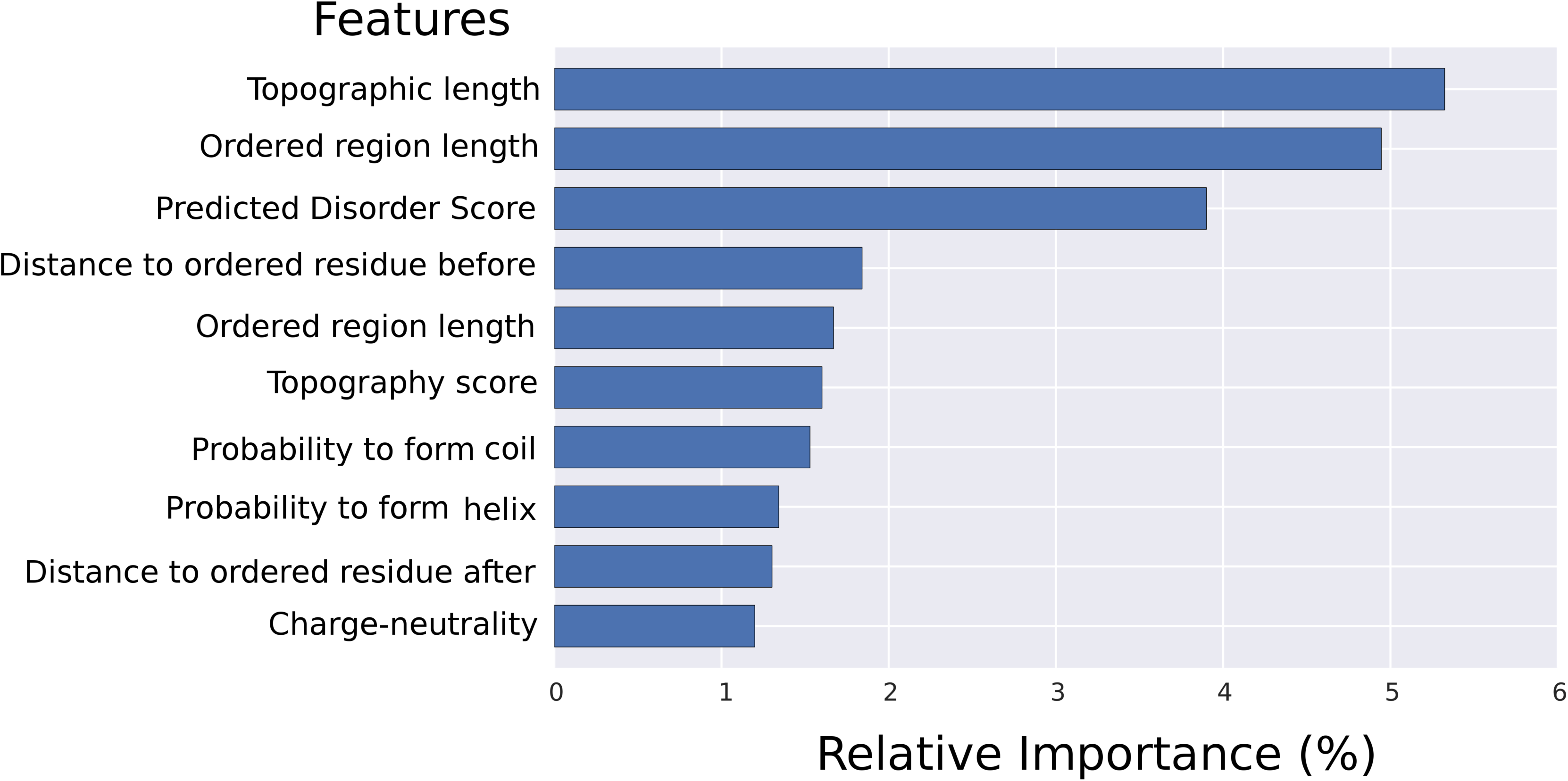
Relative feature importance. The top ten features contributing most to the prediction in the random forest features.

The other seven features in top the ten were the following. Rank 4: the relative distance to ordered residue before the current one (group 6), rank 5: length of the disordered region the current residue resides in (group 6), rank 6: the topography score (group 7), rank 7: probability of the current residue to form a coil (group 5), rank 8: probability of the current residue to form a helix (group 5), rank 9: the relative distance to ordered residue after current one (group 6), and rank 10: charge-neutrality of the current amino acid (group 4).

### True Positive Enrichment by Analyzing the Proteus Score

A common test of machine learning predictors is to analyze the true positive enrichment by constructing score plots, which is more detailed compared to recall vs. precision curves. Score plots are conventionally defined as the overlay of two independent evaluation measures, Positive Predicted Value (PPV) and recall (true positive rate) as two distinct functions of the predicted Proteus score. Ideally, both the PPV and recall should be high but there is a conflict in finding as many true positives as possible (high recall) and at the same time having a high PPV (few false positives). In reality there will always be at a trade-off between the two, which is also the main reason to use the combined measure F1. In the current case (Fig. 9A), F1 peaks at around the score of 0.5, which is also the cutoff chosen for positive prediction in the final predictor (Pcut=0.5); corresponding to 10% PPV and 23% recall as discussed above. It can be noted that after that point the PPV increases quite rapidly, and scores > 0.7 have PPV > 40%. Unfortunately there are rather few examples that obtain this high score resulting in a rather modest recall overall. Still, if the score is high we can certainly trust it to be a relatively accurate prediction. This is also reflected by analyzing the distribution of scores for protean and non-protean residues (Fig. 9B), where the score was found to be much higher for predicted protean residues than non-proteans with median values of 0.4 and 0.24 respectively, and with roughly equivalent median absolute deviations. It can also be seen that there are quite a large number of high scoring outliers in the non-protean residues. These might of course be completely wrong, but there is also a possibility that these predictions are actually sites for yet unknown interactions. Since the study of transient interaction is difficult, and the focus of the structural biology community so far has been on stable interactions that can even form crystals, there is still a lot more to be discovered if the dynamics are also taken into account.

**Fig.9.**
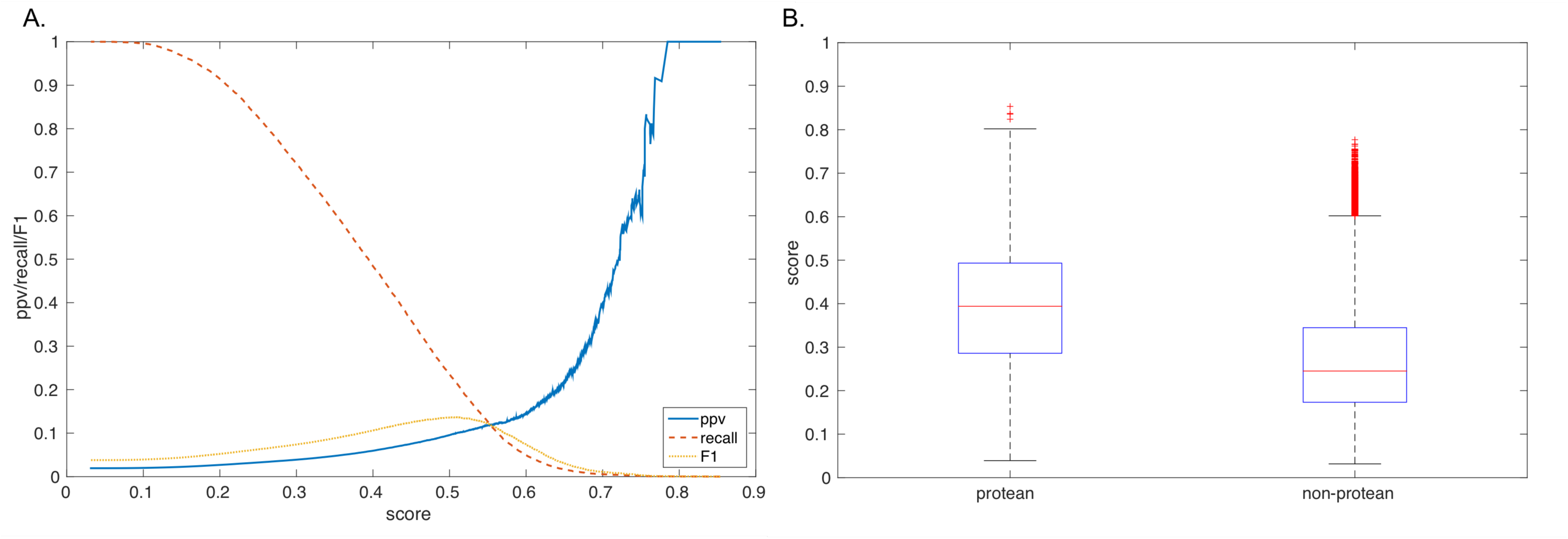
Analysis of Proteus Score for the cross-validated predictions. (**A**)Proteus score vs PPV (solid, blue), recall (dashed, red), and F1 (dotted, orange) for the cross-validated predictions. (**B**) Box plots showing the distribution of predicted Proteus scores for protean and non-protean residues. The median of the two distributions is shown by the horizontal red line in the middle of the two boxes.

### Benchmark on Independent Data Set

In any machine learning scheme it is an advantage if the final classifier can benchmarked on independent data, and against other classifiers. In the recent DISOPRED3 paper [32] the following methods were benchmarked with ANCHOR [33] MoRFpred [34], MFSPSSMpred [35], and DISOPRED3 [32] using a set of 2,209 residues out of which 163 were protean (i.e., positive examples) from nine proteins (see Methods). None of the examples in the independent set were similar to any example used in training Proteus, thus before classifying, Proteus was retrained on the full non-cross validated training set. The predictions for the other methods were generously made available by the authors of DISOPRED3 through the following link: http://bioinfadmin.cs.ucl.ac.uk/downloads/DISOPRED/suppl_data/. The evaluation measures precision, recall, F1, and MCC were calculated for all methods using the binary classification of each method (Fig. 10) and as recall vs. precision curves using the raw scores from each method (**Supplementary Fig. S8**). Overall, Proteus is better in all measures. Proteus has the highest precision (0.26 compared to 0.22 for DISOPRED3, the second best), for a much larger recall (0.56 compared to 0.28 by ANCHOR, the next best). This combined improvement in both precision and recall is also naturally reflected in a concomitant increase in the F1-score (0.35 compared to 0.18 by DISOPRED3, the next best). It also attained a higher MCC value than the other methods (0.30 compared to 0.13 by DISOPRED3). Even though the independent set is small, the high recall is particularly encouraging if Proteus is to be used as an initial step before implementing more elaborate approaches (as discussed earlier). It is crucial not to miss any true positives at an early stage.

**Fig.10.**
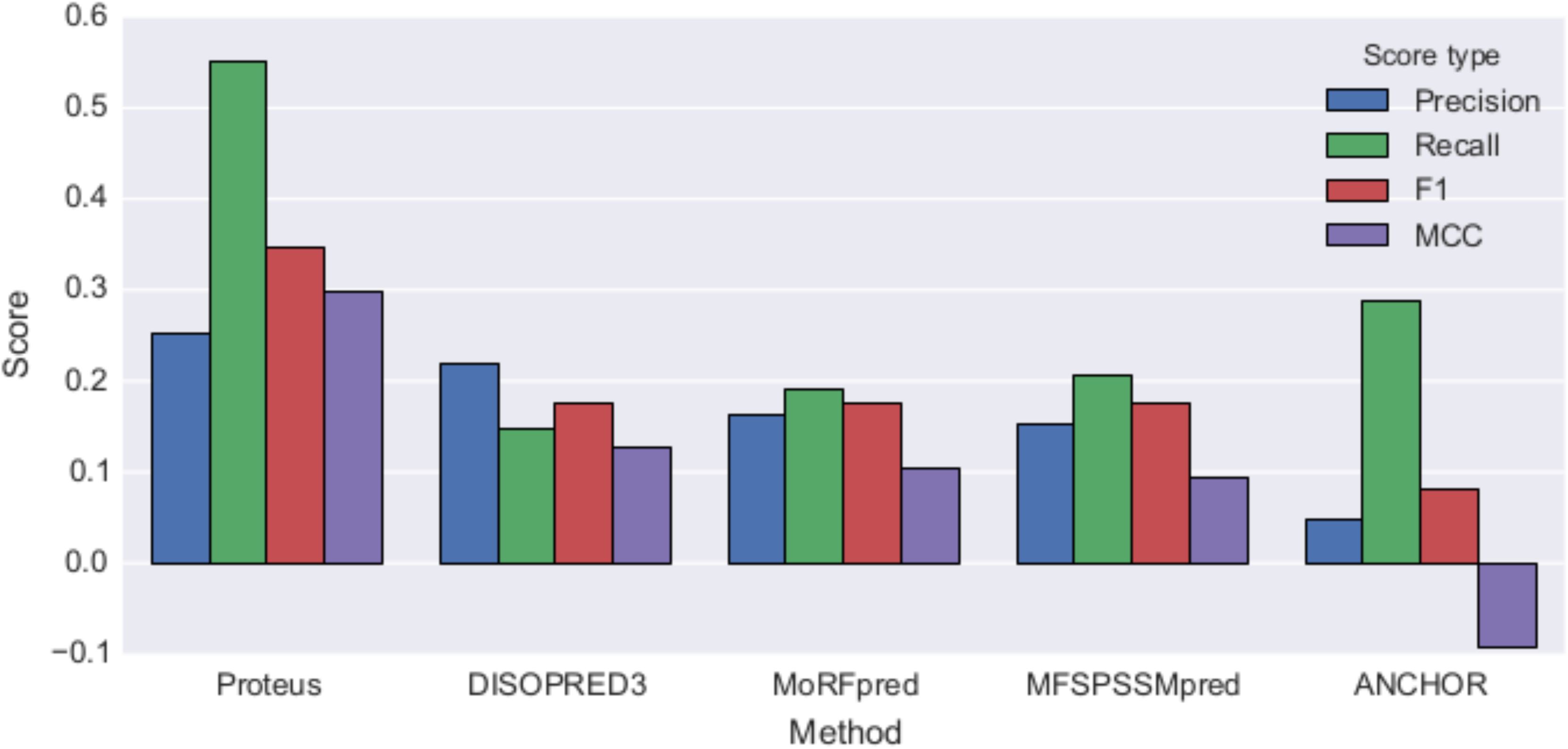
Comparison of Proteus with other classifiers using the standard evaluation measures. All methods were tested on the same validation set of nine proteins containing 2209 residues (total number of examples) with 163 protean (positive examples). Precision, Recall, F1-score and MCC tabulated for each method. Proteus predicts twice as many true positives as the second best method (55% vs. 27%) with a much higher precision.

## Conclusions

With the realization that protein disorder is involved in a range of human diseases, including cancer, cardiovascular and neurodegenerative diseases, it is important to compile more and more structural information for these proteins to understand their modus operandi. A first step in this direction is the classification and prediction of protean segments. The literature shows that there is indeed much room for improvement for the existing predictors [32]. Proteus seems to perform better than the existing predictors on the available independent dataset. Of course this has to be re-evaluated when more data becomes available. It is also possible to combine different individual methods to build hybrid methods to increase the performance even further. Given the current *state-of-the-art*, the predicted ‘protean’ segments should be considered ‘potential’ binding sites for proteins in general, whereas, for a specific interaction with known partners, the predicted segments should serve as ‘different’ starting points for model building. The built models then need to undergo stringent validation filters in an iterative cycle for screening and selection. It is also important to conceptualize the multiple sequence driven factors and realize that it is their complex coordination which holds the key consensus in promoting the ‘disorder-to-order’ transitions. The consensus is yet untangled and needs other exclusive studies to eventually be resolved, however, the current work explores certain empirically observed trend which appears to be instrumental in the transition from disorder to order. These factors include the reappearance of large hydrophobic and charged amino acids in the protean segments, which are significantly under-represented in the originally ‘disordered’ regions. The study also reflects that there is an inherent indecisiveness to adapt to a specific secondary structure (helices, strands or loops) associated with the protean segments. In other words, the protean segments remain indecisive in their choice to adapt a particular secondary structure. This is consistent with the notion of sustaining enough ‘disorder’ even in the bound form [4] which potentially helps the proteins to sustain their binding promiscuity. To conclude, the study has both a basic and an applied content, both of which should serve the IDP as well as the broad biological community.

The software package is available at https://github.com/bjornwallner/proteus

## Acknowledgments

This work was supported by grants from the Swedish Research Council (VR-NT 2012-5270), the Swedish e-Science Research Center (SeRC) and the Department of Science and Technology – Science and Engineering Research Board, India (DST-SERB research grant PDF/2015/001079). Computational resources were provided by the Swedish National Infrastructure for Computing (SNIC) at the National Supercomputing Center (NSC) in Linköping, Sweden and DST-SERB, India.

